# Centrally expressed Cav3.2 T-type calcium channel is critical for the initiation and maintenance of neuropathic pain

**DOI:** 10.1101/2022.04.27.489708

**Authors:** Sophie L. Fayad, Guillaume Ourties, Benjamin Le Gac, Baptiste Jouffre, Sylvain Lamoine, Antoine Fruquière, Sophie Laffray, Laila Gasmi, Bruno Cauli, Christophe Mallet, Emmanuel Bourinet, Thomas Bessaih, Régis C. Lambert, Nathalie Leresche

**Author notes:** Senior authors (Thomas Bessaih, Régis C. Lambert, Nathalie Leresche).

## Abstract

Cav3.2 T-type calcium channel is a major molecular actor of neuropathic pain in peripheral sensory neurons, but its involvement at the supra-spinal level is almost unknown. In the Anterior Pretectum (APT), a hub of connectivity of the somatosensory system involved in pain perception, we show that Cav3.2 channels are expressed in a sub-population of GABAergic neurons co-expressing parvalbumin (PV). In these PV-expressing neurons, Cav3.2 channels contribute to a high frequency bursting activity, which is increased in the spared nerve injury model of neuropathy. Specific deletion of Cav3.2 channels in APT neurons reduced both the initiation and maintenance of mechanical and cold allodynia. These data are a direct demonstration that centrally expressed Cav3.2 channels also play a fundamental role in pain pathophysiology.

## INTRODUCTION

Neuropathic pain is a wide spread condition with severe side effects and limited effectiveness of available treatments. Such public health issue stimulates the search for putative molecular targets, among which ion channels expressed in pain pathways are prime candidates (Bennett and Woods, 2014; Bourinet et al., 2016). In particular, at the peripheral level, it was shown that the Cav3.2 isoform of the low-threshold T-type calcium channels (T channels) are over-expressed in dorsal root ganglion (DRG) neurons after chronic constrictive injury of the sciatic nerve (Jagodic et al., 2008). Silencing the Cav3.2 isoform using intrathecal antisense oligonucleotides administration had profound anti-hyperalgesic and anti-allodynic effects in various neuropathic pain models (Bourinet et al., 2005; Messinger et al., 2009; Takahashi et al., 2010). Furthermore, specific knockout (KO) of Cav3.2 in a sub-type of DRG neurons alleviated both mechanical and cold allodynia produced by spared nerve injury (SNI) (François et al., 2015). While these data converge towards a critical role for peripherally expressed Cav3.2 in neuropathic pain, very little is known on a putative pro-nociceptive role of centrally expressed Cav3.2.

In the central nervous system, *in-situ* hybridization studies revealed strong expression of the Cav3.2 transcript in scattered neurons of the Anterior Pretectum (APT) (Talley et al., 1999), a crossroads of ascending and descending connectivity of the somatosensory system (Rees and Roberts, 1993). The idea that APT plays a key role in nociception dates back to the 1980s, when it was shown that brief low-intensity electrical or chemical stimulations of this nucleus elicited antinociception (Prado and Roberts, 1985; Rees and Roberts, 1993; Roberts and Rees, 1986). Involvement of APT in chronic pain was later confirmed by lesions and stimulations performed in different rodent pain models (Rees et al., 1995; Rossaneis and Prado, 2015; Rossaneis et al., 2014; Villarreal et al., 2003, 2004). To date the excitability of this structure has been little studied (Bokor et al., 2005) but an increase firing of fast bursting neurons was reported in a model of central pain syndrome (Murray et al., 2010). Since T channels are involved in burst generation in numerous neuronal populations (Lambert et al., 2014), we took advantage of a murine model that allows the identification of the Cav3.2 channels and their conditional deletion in specific area (François et al., 2015) to investigate their role in shaping APT neuron excitability and the initiation and maintenance of neuropathic pain.

We show that (i) Cav3.2 channels are specifically expressed in GABAergic PV-expressing neurons of the APT; (ii) bursting activity of these neurons is increased in the SNI model of neuropathic pain; (iii) Cav3.2 channels contribute to the bursting activity of the PV-expressing neurons *in vitro* and (iv) their specific deletion in the APT significantly reduces both the initiation and maintenance of neuropathic mechanical and cold allodynia in SNI mice in the spared nerve injury (SNI) model. These results are the first direct evidence that supra-spinal Cav3.2 channels play a fundamental role in pain pathophysiology.

## RESULTS

### Cav3.2 channels are expressed in PV positive (PV+) neurons of the APT

To examine Cav3.2 expression in APT neurons, we used the Cav3.2^eGFP-flox^ knock-in (KI) mouse line, in which an ecliptic GFP tag is expressed in the extracellular loop of the Cav3.2 channel (François et al., 2015). Anti-GFP labeling revealed the expression of the Cav3.2-GFP-fusion protein in a scattered subpopulation of neurons from naïve KI animals (Figure 1A, B). Co-labeling of GFP and NeuN positive cells (Figure S1) showed that 20.0±3.9% of the APT cells express the Cav3.2-GFP channel. Since a previous study performed by Bokor et al. (Bokor et al., 2005) in rats suggests that APT fast bursting neurons strongly expressed PV, the overlap between Cav3.2-GFP+ and PV+ populations was estimated using GFP and PV co-labelings (Figure S1). We thus determined that 87.1±10.0% of Cav3.2-GFP+ cells co-express PV. Conversely 91.8±5.6% of PV+ cells co-express Cav3.2-GFP confirming the overlap between the Cav3.2-GFP+ and PV+ cell populations (Figure 1C).

**Figure 1:**
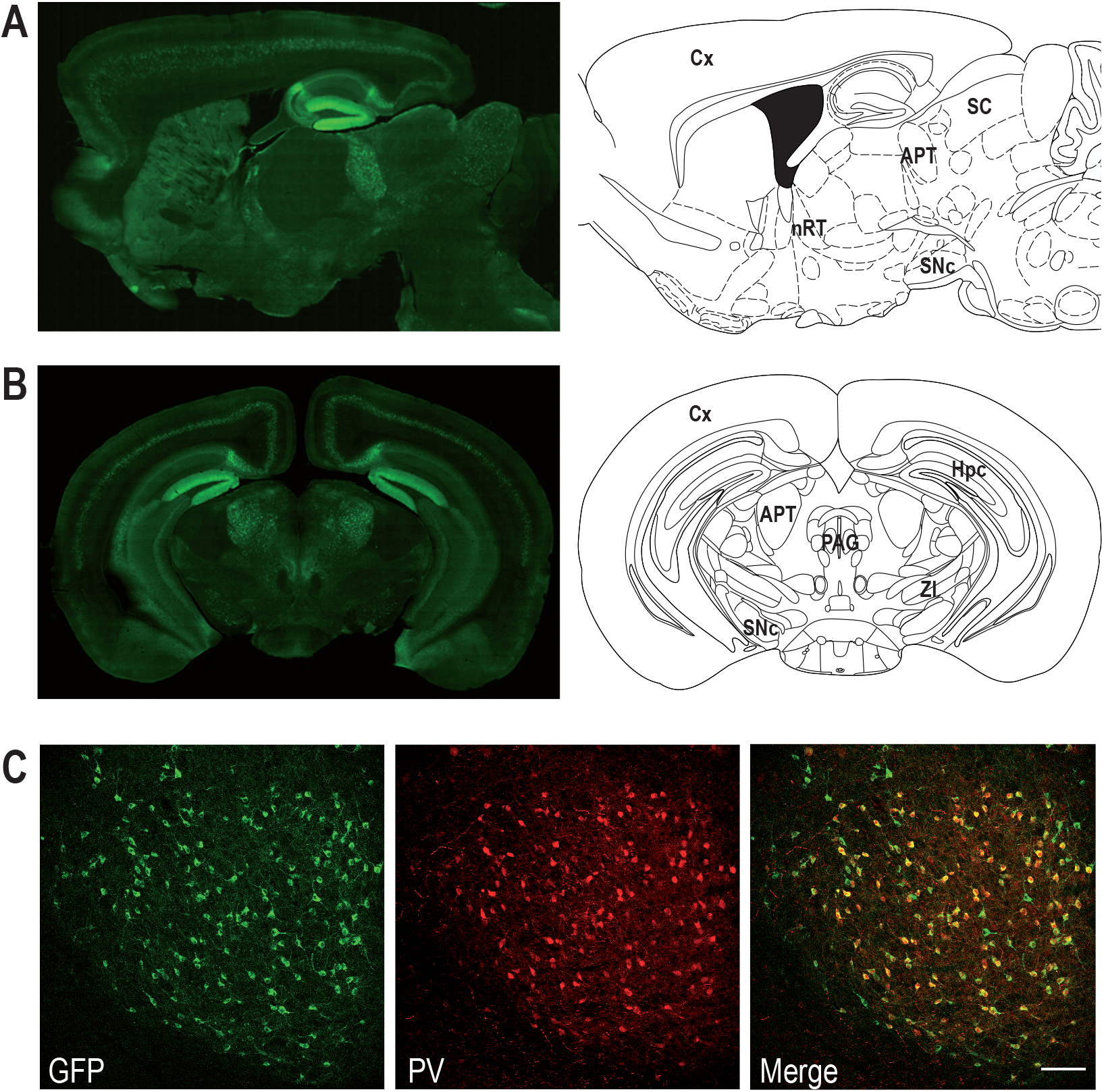
Co-expression of Cav3.2 and PV in APT neurons. (A, B) Left panels: Cav3.2-GFP immunostaining on a parasagittal (A) and a coronal (B) section of KI mice brains. Right panels: corresponding Mouse Brain Atlas slides from Paxinos and Franklin (parasagittal: 1.08mm lateral from Bregma; coronal: 2.8mm posterior from Bregma). APT: Anterior Pretectum; Cx: Cortex; Hpc: Hippocampus; nRT: Nucleus Reticularis Thalami; PAG: Periaqueductal Grey; SC: Superior Colliculi; SNc: Substantia Nigra pars compacta; ZI: Zona Incerta. (C) Confocal microscopy images of a coronal KI mouse brain section of the APT with GFP (green) and PV (red) co-labeling. Scale bar: 100 μm.

### Burst firing of PV+ neurons is increased in SNI animals

Based on this overlap, we next investigated the firing activity of this subpopulation in anaesthetized animals, using PV-Cre x Ai32 mice that selectively express channelrhodopsin-2 (ChR-2) in PV+ neurons. One to two tetrodes were lowered to the APT in the contralateral side of the SNI lesion to record multi-unit spiking activity (MUA) alongside with EEG signal (Figure 2A, B) and spike-sorting algorithms were further used to isolate one to three single-unit spiking activities per tetrode (Figure 2C). In naïve mice using Photo-assisted Identification of Neuronal Population (PINP; Lima et al., 2009), PV+ single units were identified by the reliable short-latency (5± 2.1 ms) evoked response consisting of one or more spikes elicited by each 470 nm blue-light pulse (Figure 2D). As shown in Figure 2E, 74% (n=20) of the units had a peak around ±2-3ms in their autocorrelogram, indicating their ability to elicit bursts, and are thus referred to as “bursting cells”. Their mean firing rate was 9.6±2.6Hz and bursts, consisting of 2 to 3 action potentials (mean: 2.2±0.6) occurring at 254.2±15.2Hz, represented 14.2±5.9% of the total number of spikes. The 7 remaining units showed a flat distribution of inter-spike intervals (ISI) with a mean firing rate of 4.9±1.8Hz, and are thus referred to as “regular cells” (Figure S2).

**Figure 2:**
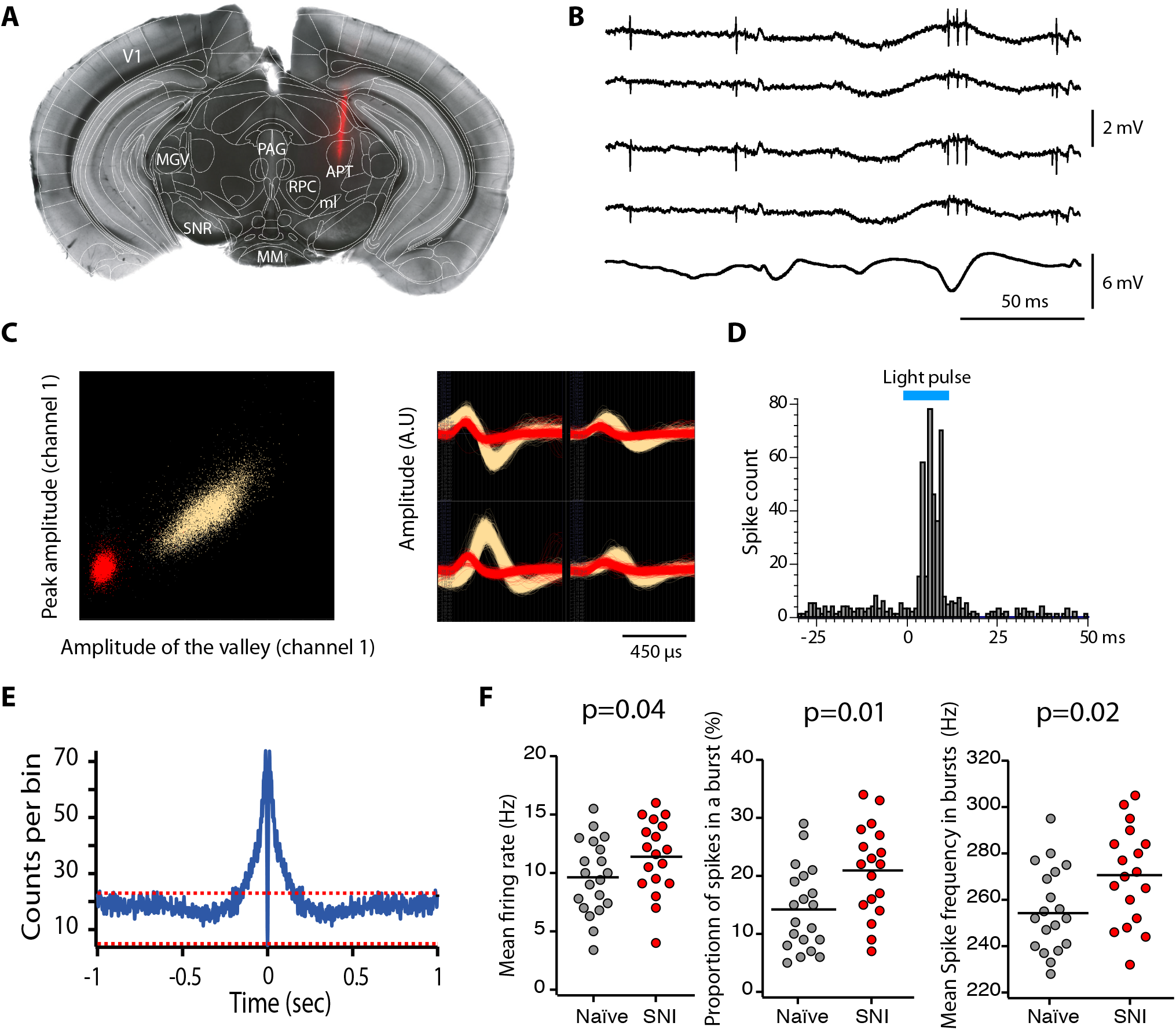
Impact of spared nerve injury in PV+ APT neurons. (A) Example image of a DiI track left by a recording electrode inserted into the APT. Red: DiI. (B) Raw signals from the four wires of a tetrode. Bottom trace shows the simultaneous EEG recording (C) Left panel: example of a tetrode recording where two units were isolated. All detected action potentials are plotted as their waveform’s amplitude from channel 1 versus the amplitude of their waveform’s valley from channel 1 (in arbitrary units (A.U.) Using this feature space arrangement, two single-unit clusters were isolated (in red and yellow). Right panel: superimposed color-coded action potential waveforms captured by each recording site of the tetrode are shown for the two identified single-units (the polarity of the signals is inverted). (D) Example of peri-stimulus time histograms illustrating spiking response to optogenetic stimulation (10ms long, represented in blue) over 100 trials of a unit categorized into the PV+ category. (E) Autocorrelogram of recorded single unit for one example cell. 1msec bins were used. Red dotted lines represent 99% confidence intervals. (F) Scatter dot plots of mean firing rate (left panel), proportion of spikes within a burst (middle panel) and mean spike frequency within a burst (right panel) for fast-bursting APT cells recorded in naïve (n=20 cells, 6 animals) and SNI (n=18 cells, 6 animals) mice.

The spiking properties of PV+ neurons were next compared between naïve and SNI animals. Although the proportion of bursting cells was similar in the two conditions (74% versus 75%), we observed a significant increase upon SNI, in the mean firing rate (Fig. 2F; left panel), in the proportion of spikes belonging to a burst (Fig. 2F; middle panel), and in the mean spike frequency within a burst (Figure 2F; right panel). No difference was observed in the mean firing rates of regular cells.

We therefore concluded that most of the PV+ and Cav3.2+ neurons of the APT are bursting neurons and that their bursting activities are enhanced in the neuropathic pain state.

### Contribution of Cav3.2 channels to PV+ APT neuron excitability

In order to estimate how Cav3.2 channels contribute to the firing activity observed *in vivo,* we characterized the excitability of the PV+ neurons using whole-cell patch-clamp recordings combined with biocytin labeling in slices obtained from PV-Cre x AI14 mice that express TdTomato in PV+ neurons. In response to a pulse of depolarizing current all APT fluorescent neurons behaved as fast spiking neurons with a high discharge rate (mean maximal firing rate upon depolarizing current of increasing amplitude: 214±112Hz, n=60, Figure 3A). In response to the injection of a hyperpolarizing current step (peak hyperpolarization ranging from −95 to - 105mV), common features were a large amplitude sag (17±6mV, n=62), indicating the presence of an Ih current, and a rebound depolarization that underlie a burst of high frequency action potentials in 57 out of 62 recorded neurons (mean number of spikes 7±4, n=57).

**Figure 3:**
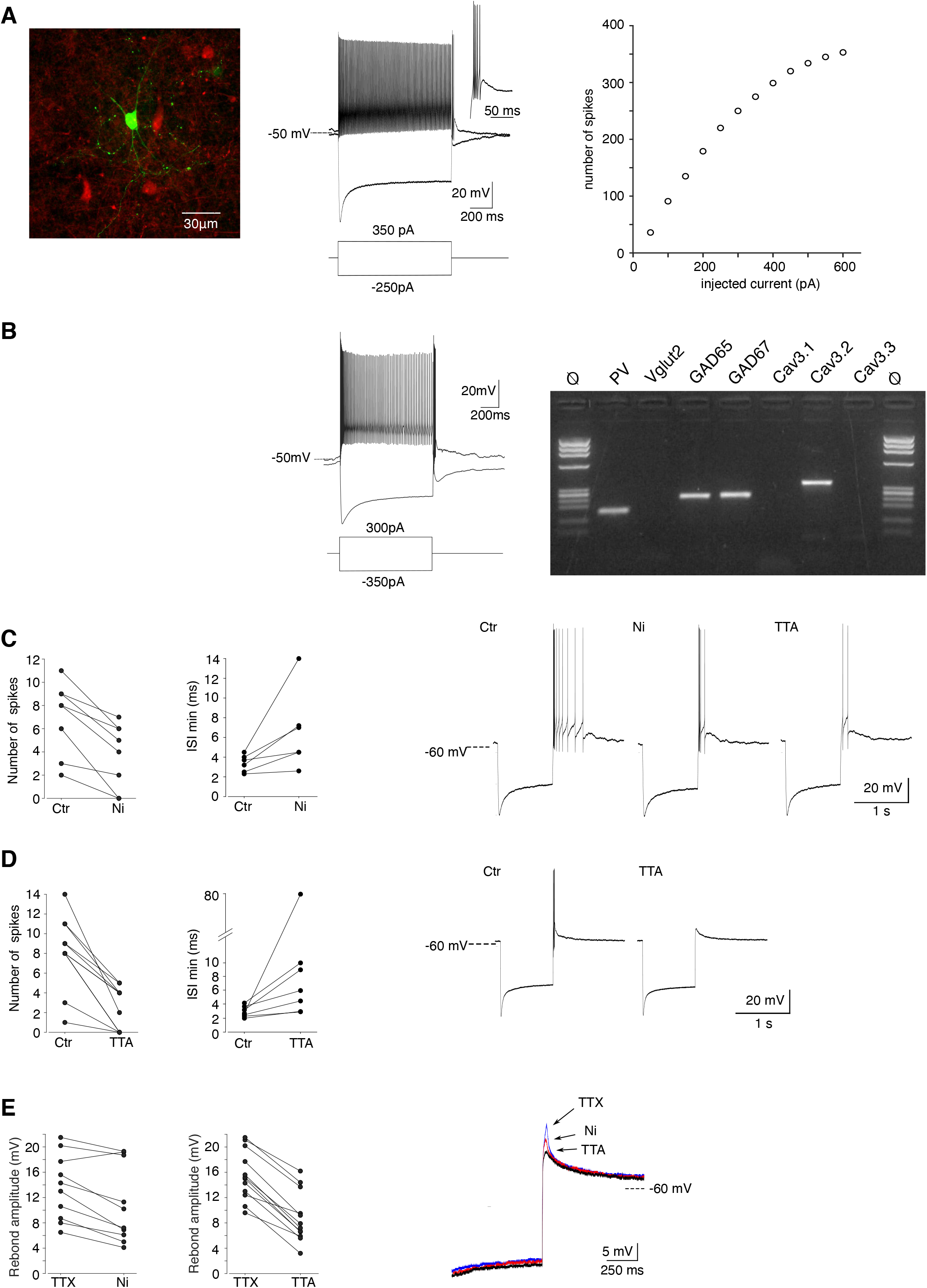
Cav3.2 channels contribute to the bursting activity of PV+ neurons. (A) Left image: recorded neuron filled with biocytin (green). Neurons in red are non-recorded PV+ neurons. Middle traces: depolarizing current injection evokes characteristic fast spiking activity. Hyperpolarizing current injection evoked a pronounced sag and a rebound high-firing burst upon repolarization. Inset: enlargement of the bursting activity. Right graph: number of spikes evoked by 1s long depolarizing current injection of increasing amplitudes. (B) Typical example of activities and scRT-PCR products observed in a PV+ neuron. Note the expression of the Cav3.2 channel in the GAD65 & 67 positive neuron. (C and D) Effect of 100μM Ni^2+^ (C) and 1μM TTA-P2 (C-D) applications on the number of spikes (left graph) and the minimal inter-spikes intervals (ISI; right graph) of the rebound bursts. Typical examples of these pharmacological effects are shown on the right. (E) Effect of 100μM Ni^2+^ (left graph) and 1μM TTAP2 (right graph) on the amplitude of the rebound depolarization observed in the presence of 0.5μM TTX. A typical example is presented in superimposed traces presented on the right.

Transcriptomic characterization of PV+ neurons was further performed by combining patch-clamp recordings and multiplex single-cell reverse transcriptase PCR (scRT-PCR; Figure 3B). All neurons expressed PV transcript and among them, 73.9% expressed GAD65 and/or GAD67 mRNA (n=17/23), the remaining ones (n=6/23) expressed Vglut2 mRNA. Importantly the Cav3.2 transcript was present in 13 out of the 17 GABAergic neurons but never detected in glutamatergic neurons. mRNA of the two other Cav3 isoforms was also detected, albeit less frequently (Cav3.1: n=10/13; Cav3.3: n=3/13) in GABAergic neurons co-expressing Cav3.2 and in glutamatergic neurons (Cav3.1: n=5/6; Cav3.3: n=1/6). These results strongly suggest that PV+ neurons expressing the Cav3.2 channels are fast spiking GABAergic neurons.

We then investigated whether the Cav3.2 isoform contributed to the rebound depolarization and its associated high frequency burst firing. As shown in Figures 3C and 3E, application of 100μM Ni^2+^, a concentration that specifically blocks Cav3.2 channels (Lee et al., 1999), significantly decreased the number of action potentials (ctr: 7.0±3.1 / Ni^2+^: 3.8±2.8; n=8; p=0.0039), the firing frequency (min ISI: ctr: 3.4±0.9ms / Ni^2+^: 6.6±4.0ms; n=6; p=0.0156) and the amplitude of the underlying depolarization measured in the presence of TTX (ctr: 13.6±5.2/Ni^2+^: 10.8±6.1mV; n=10; p=0.0029). A block of all T-channels isoforms using the pan antagonist TTA-P2 produced an even stronger decrease of the rebound activity, as expected from the results of the scRT-PCR (Figure 2D, E).

### Impact of Cav3.2 channel deletion in APT on mechanical and cold sensitivity

The Cav3.2^eGFP-flox^ KI mouse line not only allows the visualization of Cav3.2 channels but also their deletion (Figure S4). Mechanical and cold allodynia, the cardinal somatosensory phenotypes typical of the SNI model were thus compared between mice that were bilaterally injected in the APT with either an AAV-Cre-mCherry virus (APT-KO mice; n=6 males and 6 females) or an AAV-mCherry (control KI mice; n=7 males and 9 females; Figure 4A) and then two weeks later subjected to the SNI surgery to induce the neuropathy. Mechanical sensitivity was assessed by measuring Paw Withdrawal Threshold (PWT) on the operated hindpaw using Von Frey filaments (Chaplan et al., 1994). Prior to SNI procedure, 2 baseline measurements were performed, before and 2 weeks after viral injections. No significant differences were found in the PWT between these 2 measurements within each group or between KI and APT-KO mice, indicating that neither the viral injection by itself, nor the local deletion of Cav3.2 impacts acute mechanical sensitivity. Baseline measurements were therefore averaged within each group (Figure 4B). A statistically significant decrease in PWT was observed in KI mice following SNI from day 14 post-surgery, confirming the development of a mechanical allodynia. Importantly, this decrease was strongly attenuated in APT-KO compared to KI mice (Figures 4B and S4B). This difference between APT-KO mice and their KI littermates was not due to motor or coordination deficits (Figure S4C, S4D).

**Figure 4:**
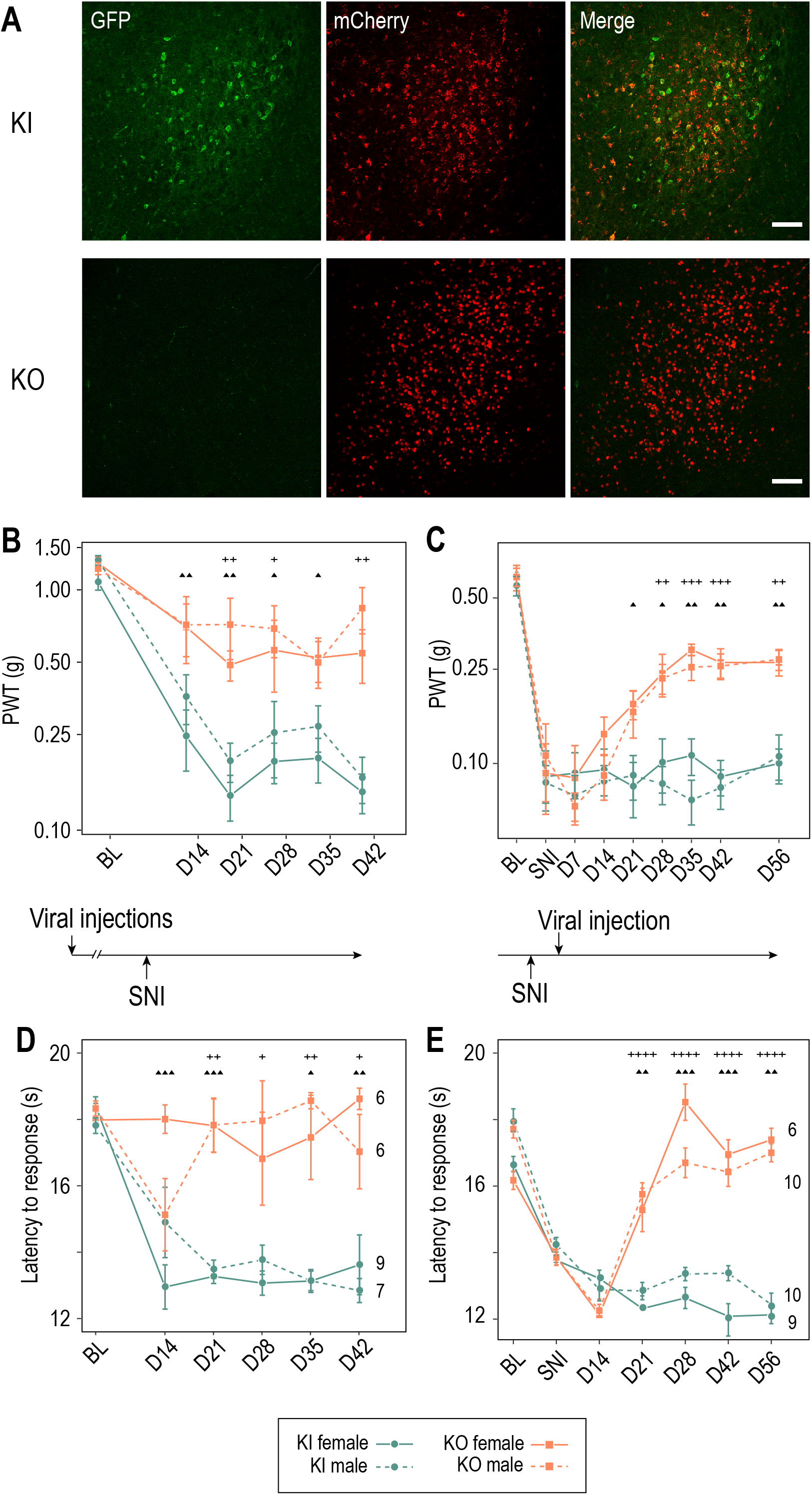
Cav3.2 preventive and curative knock-out in the APT alleviates neuropathy induced by SNI. (A) Confocal microscopy images of GFP (green) and mCherry (red) co-labeling in the APT of KI-Cav3.2-GFP (Control) mice and KO-Cav3.2-APT (KO) bilaterally injected in the APT with AAV8-hSyn-mCherry and AAV8-hSyn-mCherry-CRE virus respectively, and further tested for mechanical and cold sensitivity. Scale bar: 100μm. (B - E) Neuropathic behaviors were tested in male (dashed lines) and female (solid lines) mice with preventive (B, D) and curative (C, E) KO of Cav3.2 in the APT (orange, ■), and in control KI mice (green, •). (B, C) Mechanical sensitivity was assessed by measuring Paw withdrawal thresholds (PWT) in response to Von Frey filaments stimulations using the up-and-down method. (D, E) Cold sensitivity was assessed by measuring the paw withdrawal latency in response to immersion in 18°C water. For preventive KO (B, D), mice were tested prior to the SNI (BL) and once a week during the 6 subsequent weeks (days 14 to 48). For curative KO (C, E), mechanical and cold sensitivity were assessed before (BL) and 14 days after SNI (SNI), and tested during several weeks following the subsequent viral injections (days 14 to 56 after viral injection). Significant differences: *p < 0.05 **, p < 0.01 ***, p < 0.001 ****, p < 0.0001 (males: +; females: ▲).

Cold sensitivity was also assessed using the paw immersion test (Figure 4D). While KI mice showed a decrease in withdraw latency after SNI when the operated paw was immersed in a water bath at 18°, indicating the development of a cold allodynia, this phenotype was abolished in the preventive APT-KO mice. Altogether, these results clearly indicate that APT Cav3.2 channels contribute to the initiation of mechanical and cold allodynia characterizing the SNI model in both male and female mice.

We next investigated whether these allodynic SNI features could be reversed by a curative KO strategy of APT-Cav3.2. We thus injected the Cre-expressing virus 2 weeks after surgery, when allodynic symptoms reach their peak. Both mechanical and cold allodynia were rapidly rescued since tactile threshold lowering was drastically reduced (Figure 4C) and paw withdrawal latencies returned to preoperative baseline values (Figure 4E).

Altogether, these results show that Cav3.2 expressed in the APT is not only involved in the initiation but also in the maintenance of neuropathic phenotype in the SNI model.

## DISCUSSION

We showed that deletion of Cav3.2 channels expressed in a GABAergic and PV expressing sub-population of APT neurons leads to an anti-allodynic effect in the SNI neuropathic pain model.

Our *in vitro* data indicate that 92% of APT-PV+ neurons are able to discharge bursts of action potentials at high frequency underpinned by a large transient depolarization due to the activation of T channels. In contrast to the well-recognized role played by the Cav3.1 and 3.3 isoforms of the T channels (Kim et al., 2001; Lee et al., 2014; Pellegrini et al., 2016), little evidence exist for the involvement of the Cav3.2 isoform in burst generation in the CNS (Candelas et al., 2019; Dumenieu et al., 2018). Here, the specific blockade of this isoform greatly reduced the amplitude of the depolarization underlying burst generation and decreased the number and frequency of action potentials within a burst. Therefore, although our transcriptomic and pharmacological results suggest that APT-PV+ GABAergic neurons express multiple isoforms of the T channels, we demonstrate that the Cav3.2 isoform has an essential contribution to the bursting behavior in this sub-population of APT neurons.

Available data about the excitability of APT neurons are very limited. Bokor et al. (Bokor et al., 2005) reported a heterogeneous firing pattern in rat APT neurons with about 25% of the them presenting high-frequency discharges. Although tested on only 3 neurons, these fast-bursting neurons were described as strongly expressing PV. Deciphering the roles of a structure that displays such heterogeneity clearly requires recordings of identified neuronal subpopulation. The use of the PINP method allowed us to perform such specific characterization and our *in vivo* recordings of APT-PV+ neurons showed that 74% of these neurons discharge in bursts with a clear enhancement of bursting activity in SNI mice. Investigating changes in neuronal spiking activity in spinal-lesioned rats, Murray et al. (Murray et al., 2010) also described an increased firing rate of APT neurons with a higher percentage of neurons exhibiting at least one burst compared with sham-operated controls. The increase in bursting behavior reported here were more pronounced than in Murray et al. (Murray et al., 2010) as not only the number of bursts but also the number of spikes and their frequency were enhanced indicating a profound change in this class of APT neurons in neuropathic condition. This disparity likely stems from our ability to record an identified subpopulation of APT neurons, although a difference between neuropathic models cannot be excluded. When considering other brain structures, modification of high-frequency bursting activities in pain condition have been reported in human, as well as in various experimental models, and this is particularly well documented in the thalamocortical system. In awake patients with neurogenic pain, recordings from thalamic neurons show an increase in the occurrence of high-frequency bursts (Jeanmonod et al., 1996; Lenz et al., 1989; Llinas et al., 1999). Similar higher proportions of bursting neurons have been reported in ventralis postero-lateralis thalamic neurons following contusive spinal cord lesions in rats (Gerke et al., 2003) and monkeys (Weng et al., 2000). Furthermore, in the same thalamic nucleus, an increase in the frequency of T channel dependent bursts was reported following induction of visceral pain in mice (Kim et al., 2003). The increased burst firing in APT-PV+ neurons in neuropathic condition prompted us to investigate whether Cav3.2 channels may contribute to this phenotype. Our behavioral analysis firstly showed that removing Cav3.2 specifically from APT-PV+ neurons strongly reduced mechanical and cold allodynia evoked by SNI. Furthermore, removing the Cav3.2 channels when the SNI effects were fully developed rescues the different allodynic symptoms indicating that Cav3.2 channels not only contribute to the development but also to the maintenance of the pain phenotype and strongly suggests their direct electrogenic implication in sustaining allodynia. Since these effects were observed on both male and female mice, the Cav3.2 dependent modification in APT-PV+ neurons activities sustaining allodynia is not sex-specific, unlike what was recently reported for layer 5 PV+ neurons in the prelimbic region of the medial prefrontal cortex using the same neuropathic model (Jones and Sheets, 2020). Such lack of sex specificity suggests that this mechanism is a fundamental component of chronic pain processes.

Although multiple data point towards Cav3.2 as a main actor in pain processing at the peripheral level (Bourinet et al., 2005, 2016; François et al., 2015; Jagodic et al., 2008; Messinger et al., 2009; Takahashi et al., 2010), very little is known at supra-spinal levels. Intracerebroventricular injection of the specific T channel antagonist, TTA-A2, induced analgesia blunting the effect of paracetamol, the most widely used remedy to treat mild pain, an effect abolished in Cav3.2^-/-^ mice (Kerckhove et al., 2014). Similarly, using various less specific T channel antagonist, Chen et al. (Chen et al., 2010) showed a T channel dependent activation of ERK in the paraventricular thalamus following acid-induced chronic muscle pain that was also abolished in Cav3.2^-/-^ mice. However, such experiments only suggested a central pronociceptive role of Cav3.2 channels. Our results, showing that local deletion of Cav3.2 channels in 20% of the APT neurons drastically reduced allodynia, provide the first direct evidence of the involvement of central Cav3.2 channels in neuropathic pain.

From a functional point of view, as APT-PV+ neurons project to high-order somatosensory thalamic nucleic (Bokor et al., 2005; Giber et al., 2008), we may hypothesize that this anti-allodynic effect is the consequence of a direct implication of this neuronal population in the processing of nociceptive afferences from the periphery. However, It remains to be determined whether the APT-PV+ neurons may also contribute to the descending control of pain perception as anatomical investigation of the APT described descending pathways impinging through complex routes on various spinal cord neuronal types (Genaro et al., 2019; Rees and Roberts, 1993; Terenzi et al., 1991, 1992; Villarreal et al., 2004).

In conclusion, our data point to Cav3.2 channels as a prime target for developing innovative analgesic pharmacology that will not only act at the peripheral level but also in central structures. Furthermore, they highlight the need to study the functional expression of channels in specific neuronal population to decipher the complex pain processing that occurs in the central nervous system.

## Acknowledgments

This work was supported by operating grants from the Agence Nationale de la Recherche (ANR Pain-T and “Investissements d’Avenir” program I-Site CAP 20-25).

## Author Contributions

S.L.F., G.O., B.L.G., B.J., S.L., A.F., S.L., L.G., T.B., R.C.L and N.L. conducted the experiments; M.C., M.A.L., S.L.F., G.O., B.L.G., B.C., C.M, T.B; R.C.L. and N.L. analyzed the experiments; S.L.F., T.B., R.C.L., and N.L. wrote the article; T.B., E.B., R.C.L. and N.L. directed the work.

---

**Supplement to Figure 1:**
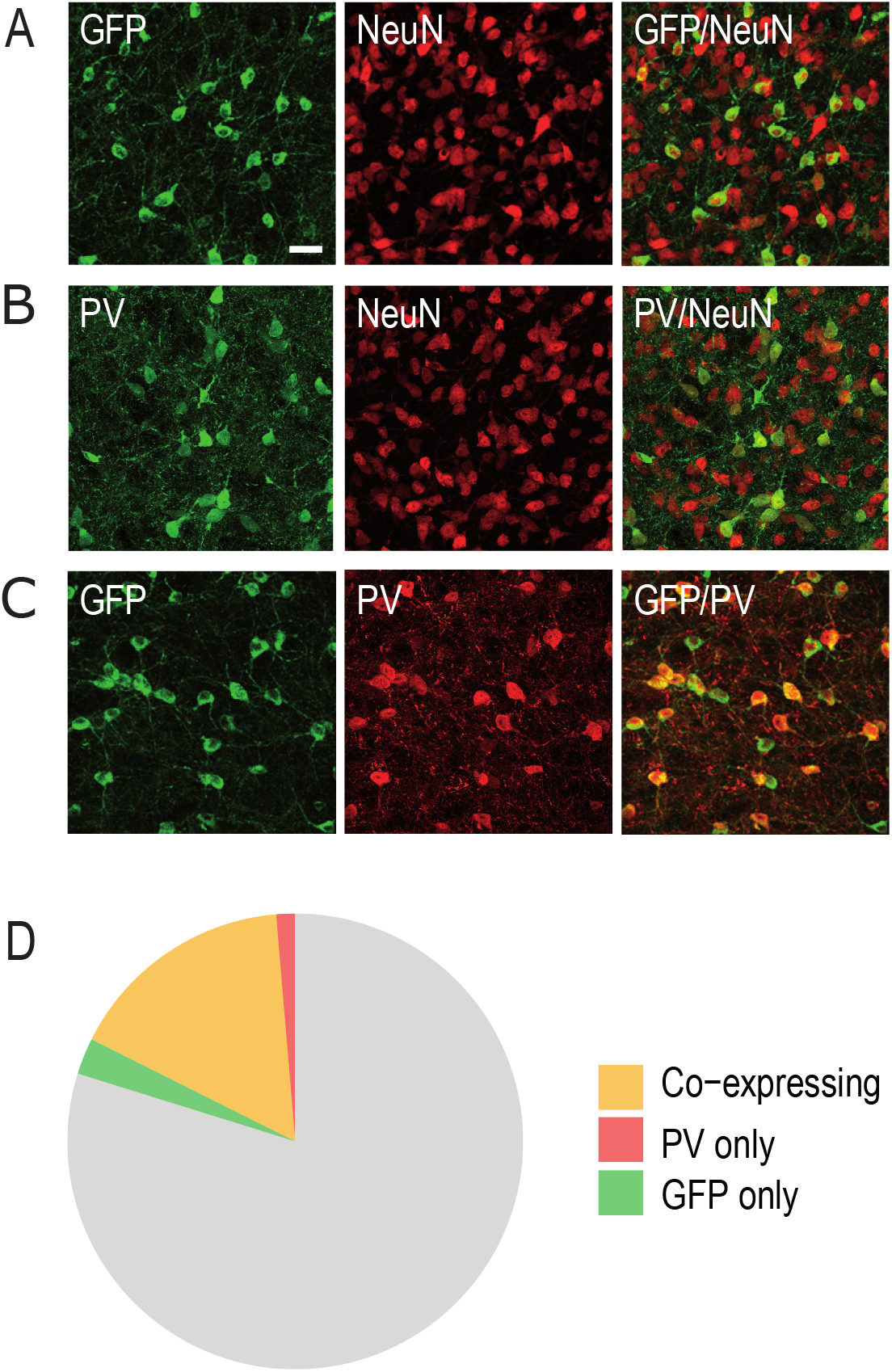
APT neurons expressing GFP and PV. (A-C) Confocal microscopy images of the co-labelings performed in the APT to estimate the proportion of GFP neurons (A: GFP in green, NeuN in red) and PV neurons (B: PV in green, NeuN in red) and the overlap between these populations (C: GFP in green, PV in red). Scale bar: 30μm. (D) Diagram representing the percentage of neurons expressing GFP and PV in the APT.

**Supplement to Figure 2:**
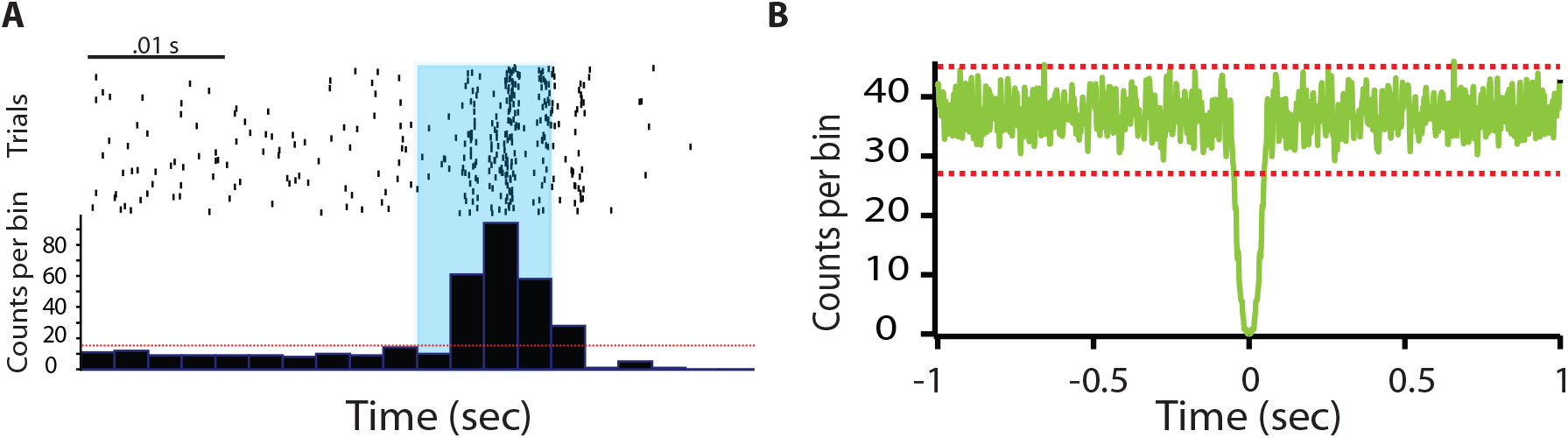
APT regular spiking neurons. A) Example of peri-stimulus time histograms illustrating spiking response to optogenetic stimulation (10ms long, represented in blue) over 100 trials of a unit categorized into the PV+ category. (B) Autocorrelogram of recorded single unit for one example cells. 1msec bins were used. Red dotted lines represent 99% confidence intervals.

**Supplement to Fig 4:**
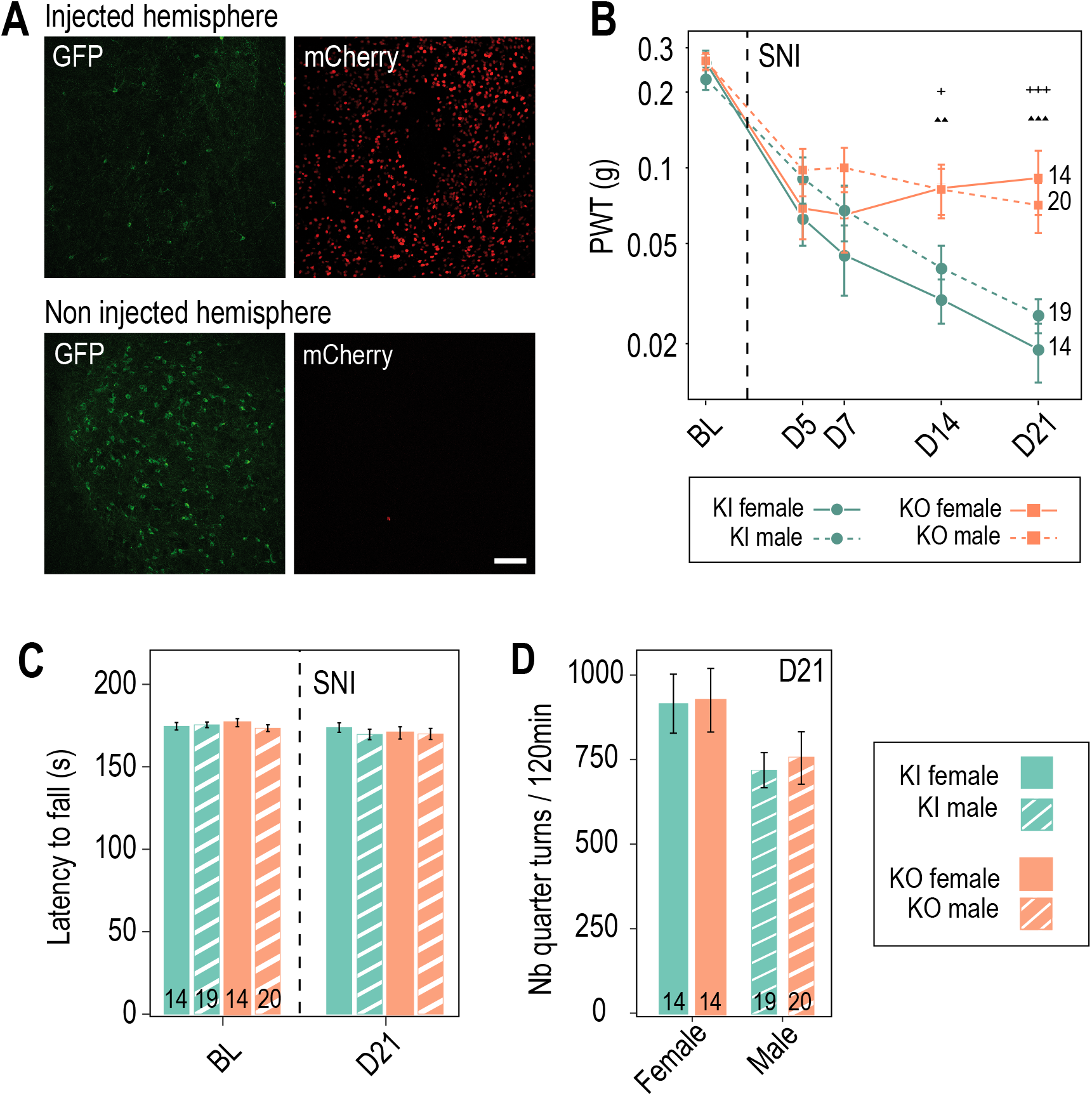
Preventive Cav3.2 knock-out in the APT has no impact on motor coordination and spontaneous locomotion. (A) Confocal microscopy images of GFP-labeling (green) and mCherry expression (red) in the APT of a KI mouse unilaterally injected with AAV8-hSyn-Cre-mCherry virus (up panel: injected hemisphere; down panel: non-injected hemisphere). Scale bar: 100μm. Note the drastic reduction in the number of GFP+ neurons observed two weeks post viral injection in the injected hemisphere. (B-D) Motor coordination and spontaneous locomotion assessed in control KI mice (green, •) and mice with preventive APT Cav3.2 KO (orange, ■) subjected to SNI. (B) Control of the development of mechanical allodynia in male (dashed lines) and female (solid lines) mice. Note that as for the cohorts presented in Figure 4, preventive Cav3.2 KO reduced allodynia. Significant differences: *p < 0.05 **, p < 0.01 ***, p < 0.001 ****, p < 0.0001 (males: +; females: ▲) (C) Latency to fall from an accelerating rotarod wheel (0-16rpm over 3min) was measured in males (dashed bars) and females (solid bars), before (BL) and after (day 21) SNI surgery. (D) Number of quarter turns were detected in a circular corridor over 120min, measured in males (dashed bars) and females (solid bars), 21 days after SNI surgery.

## Materials and methods

### LEAD CONTACT AND MATERIALS AVAILABILITY

For each figure, data are included in the “Figure – source data” files.

Further information and requests for resources and reagents should be directed to and will be fulfilled by the Lead Contact, Régis C Lambert

### EXPERIMENTAL MODEL AND SUBJECT DETAILS

#### Animals

Cav3.2^eGFP-flox^ (Cacna1h^tm1.1(epH)Ebou^/J) knock in (KI) mouse line was obtained from Dr. E. Bourinet (François et al., 2015). Briefly, these mice express the ecliptic GFP (eGFP) in an extracellular loop of the Cav3.2 T-channel. LoxP sites were inserted allowing the deletion of the Cav3.2-eGFP coding sequence by *cre*-recombinase (*cre*). Mouse lines of Parvalbumin (PV)*cre* (B6.129P2-Pvalb^tm1(cre)Arbr^/J, stock #017320) mice expressing *cre* under the parvalbumin promoter, Ai14 (B6.Cg-Gt(ROSA)26Sor^tm14(CAG-tdTomato)Hze^/J, stock #007914) mice expressing tdTomato reporter, and Ai32 (B6.Cg-Gt(ROSA)26Sor^tm32(CAG-COPC4*H134E/EYPF)Hze/^J, stock #024109) mice expressing channelrhodopsin-2 (ChR2)-eYFP were purchased from Jackson lab. PV*cre*-Ai14 and PVcre-Ai32 mice were obtained by crossing female PV*cre* with male Ai14 or Ai32 mice, respectively. Animals were housed in groups of 2 to 5 per cage, with a 12/12 hours light/dark cycle in a pathogen-free facility maintained at 22–24°C, and access to food and water *ad libidum.* All procedures complied with the ethical guidelines of the Federation for Laboratory Animal Science Associations (FELASA) and with the approval of the French National Consultative Ethics Committee for health and life sciences (authorization number: 17958).

### METHOD DETAILS

#### Immunocytochemistry and imaging

Mice (P28 to P217) were anesthetized with 2% isoflurane, injected with a lethal dose of pentobarbital (150 mg/kg) and transcardiacally perfused with a 4°C solution of Artificial Cerebro-Spinal Fluid (ACSF), oxygenated with a mixture 95%O2 / 5%CO2, containing (in mM): 125 NaCl, 2.5 KCl, 2 CaCl2, 1 MgCl2, 1.25 NaH2PO4, 26 NaHCO3, 25 glucose. The brains were then rapidly extracted from the skull and incubated in a solution of paraformaldehyde (PFA) diluted to 4% in phosphate buffer solution (PBS: Phosphate Buffer Saline) for post-fixation overnight at 4°C. 40μm thick coronal slices were cut using a vibratome (Leica VT1000S) in PBS.

All immunohistochemical staining were performed at room temperature. The slices were rinsed three times in Tris Buffer Saline (TBS, containing 50mM Tris Base and 150mM NaCl) at pH 8.4, then incubated in TBS solution supplemented with 0.05% Tween20, 0.2% Triton100X and 10% Donkey Serum (TBSTD) for 1h30. The slices were then incubated with the primary antibodies (see Table) in TBSTD solution overnight, rinsed 3 times with TBS, incubated with secondary antibodies (see Table) in TBSTD for 2h and rinsed again 3 times before mounting between slide and coverslip in Fluoromount.

Whole slice images were acquired by epifluorescence microscopy under a macroscope (Axio Zoom, V16 Zeiss). Mosaics were made at 200X magnification and processed with the ZEN software (Zeiss). The images used to study the co-expressions were acquired with confocal microscopy (Leica TCS SP5) at 20X and 63X objectives.

#### Spared nerve injury model

Spared nerve injury (SNI) was performed as previously described (Decosterd and Woolf, 2000) under ketamine-xylasine anesthesia (100mg/kg and 10 mg/kg respectively) on male and female KI mice (10 to 15 weeks old). The left thigh to be operated on was slightly elevated and an incision of about 1cm is made between the hip and the knee. The muscles enclosing the sciatic nerve compartment were moved apart with a round-ended bent scissor. The common peroneal and tibial branches of the sciatic nerve were exposed and tightly ligated with 6-0 silk suture. A fragment of nerve was transected distally to the ligation. The sural branch was left intact. The muscles were then put back in apposition and the skin was sutured using 4-0 Vicryl (Ethicon). Mice were then kept in a 37°C warming chamber until recovery from anesthesia, before being returned to their homecages.

#### Surgery and preparation for in vivo electrophysiological recordings

Mice (males; 8 to 12 weeks; Naïve mice n=5; SNI mice n=4) were anesthetized with isoflurane vaporized in a mixture of oxygen and air (4% for induction, 1.5 to 2% during surgery). Body temperature was maintained at 37°C via a servo-controlled heating blanket and a rectal thermometer (Harvard Apparatus, Holliston, MA). Bupivacaine (subcutaneous) was administered in the regions to be incised 15 min prior to the first incision. Mice were placed in a stereotaxic apparatus and a craniotomy was made directly above the anterior pretectal nucleus (−2.7 to −3.0 mm A/P, 1 to 1.2 mm M/L). To minimize damage during electrode penetration, the dura was resected and the exposed surface was coated with a layer of silicon oil. After electrodes placement, isoflurane was decreased to 0.8 to 1 %. If the animals presented any sign of discomfort, the percentage of isoflurane was increased. Electrodes were lowered to the anterior pretectal nucleus based on readings from the micromanipulator (depths: 2600 to 2900 μm).

#### In vivo electrophysiological recordings

Recordings of units were obtained using quartz-insulated platinum/tungsten (90%/10%) tetrodes (~1–2 MOhm, Thomas Recording). Before insertion, the rear of the tetrodes was painted with fluorescent 1,1’-dioctadecyl-3,3,3’,3’-tetramethyl indocarbocyanine perchlorate (DiI, 10% in ethanol, Invitrogen). As this dye is a lipophilic neuronal tracer, it allowed assessment of the recording depth. 1 to 3 tetrodes were guided independently at 1 μm resolution through a five-channel concentric microdrive head (Head05-cube-305-305-b, Thomas Recording Gmbh) with 305 μm inter-electrode spacing.

Raw signals were filtered (600–6000 Hz; Neuralynx recording systems), amplified (5000x), digitized at 33,657 Hz, and stored with stimulus markers (Cheetah 5 software; Neuralynx). Waveforms crossing set thresholds (300-500 μV) were captured via the A/D card and analyzed off-line. At the end of the experiment, mice were injected with a lethal dose of euthasol, and the brains were removed and placed in a 4% paraformaldehyde solution for 48 h. The brains were then transferred and stored in phosphate buffer saline (0.1 M). Sections (280 μm thick) containing the anterior pretectal nucleus were cut with a vibratome (Leica VT1000S). Sections were then mounted in Fluoromount medium.

#### Slice preparation

13 male and 11 female PVcre-AI14 mice (P23 to P123, median: P35) were anesthetized with isoflurane before decapitation. The brain was carefully removed and placed for a few minutes into a 4-8°C bicarbonate-buffered saline (BBS) solution containing (in mM): 125 NaCl, 2.5 KCl, 2 CaCl2, 1 MgCl2, 1.25 NaH2PO4, 26 NaHCO3 and 25 glucose (osmolarity: 305 mOsm; pH 7.3 after equilibration with 95% O2 and 5% CO2). Coronal slices (250 μm) were then cut using a vibratome (Campden Instruments 700 SMZ2). The slicing procedure was performed in an ice-cold solution containing (in mM): 130 potassium gluconate, 15 KCl, 2 EGTA, 20 HEPES, 25 glucose, 1 CaCl2 and 6 MgCl2 (304mOsm, pH 7.4 after equilibration). Slices were then transferred for few minutes to a solution containing (in mM): 225 D-mannitol, 2.5 KCl, 1.25 NaH2PO4, 25 NaHCO3, 25 glucose, 1 CaCl2 and 6 MgCl2 (310 mOsm, 4-8°C, oxygenated with 95% O2 / 5%CO2), and finally stored for the rest of the experiment at 32°C in oxygenated BBS. For all recordings, slices were continuously perfused with oxygenated BBS at 32°C.

#### In vitro whole-cell patch-clamp recording

Brain slices were screened for fluorescent neurons using a filter set that allowed us to detect dt-tomato fluorescence. Neurons were visualized and patched with borosilicate pipettes (resistance 3–5 MOhm). The intracellular solution contained (in mM): 140 potassium gluconate, 3 MgCl2, 10 HEPES, 0.2 EGTA, 4 disodium ATP (pH 7.3; 300 mOsm). For some experiments, biocytine (2mg/ml) was added to the intracellular solution. Patch-clamp electrodes were connected to a AxoPatch 200B (Axon Instrument) amplifier. Protocols and acquisitions were controlled by the Clampex software (Molecular Devices). The membrane potentials were filtered by a 4-pole Bessel filter set at a corner frequency of 2 kHz and digitized on-line at a sampling rate of 20 kHz. The access resistance was 10-20 MOhm and was monitored throughout the experiment. Data were discarded if the access resistance changed by more than 15% during the experiment. Current clamp recordings were performed in the continuous presence of 10μM CNQX and 1μM SR95531 to suppress spontaneous synaptic activities.

At the end of the recordings, slices containing biocytin-filled neurons were fixed overnight by immersion in paraformaldehyde (4% in PBS 1M) and then washed with 1M Phosphate Buffer saline (PBS). After incubating the slices with Triton (0.2% in PBS 1M) for 1h, biocytine filled neurons were revealed using Streptavidin Alexa Fluor 488 (1:1000; 3 h in dark; Invitrogen). Slices were then washed in PBS (1 h) before being mounted on cover slides.

#### Cytoplasm harvesting and scRT-PCR

For ScRT-PCR, recordings were performed on slices obtained from 4 male and 2 female PVcre-AI14 mice (P15 to P21). At the end of the whole-cell recording, lasting less than 15 min, the cytoplasmic content was aspirated in the recording pipette. The pipette’s content was expelled into a test tube and reverse transcription (RT) was performed in a final volume of 10 μl, as described previously (Lambolez et al., 1992). The scRT-PCR protocol was designed to probe simultaneously the expression of Cav3 isotypes, GAD65/67, VGluT2 and PV. Two-steps amplification was performed essentially as described (Cauli et al., 1997; Devienne et al., 2018). Briefly, cDNAs present in the 10 μl reverse transcription reaction were first amplified simultaneously using all external primer pairs listed in the Key Ressources Table. Taq polymerase and 20 pmol of each primer were added to the buffer supplied by the manufacturer (final volume, 100 μl), and 20 cycles (94°C, 30 s; 60°C, 30 s; 72°C, 35 s) of PCR were run. Second rounds of PCR were performed using 1 μl of the first PCR product as a template. In this second round, each cDNA was amplified individually using its specific nested primer pair (Key Ressources Table) by performing 35 PCR cycles (as described above). 10 μl of each individual PCR product were run on a 2 % agarose gel stained with ethidium bromide using ΦX174 digested by *HaeIII* as a molecular weight marker.

#### Virus stereotaxic injections

Male and female KI (6 to 8 weeks) were anesthetized with a ketamine-xylazine mixture (100mg/kg and 10mg/kg, respectively) and placed on a heating pad. A vitamin B12 eye drop (Twelve TVM) was applied to the eyes and a subcutaneous injection of sterile saline solution (NaCl 0.9%) was performed to prevent dry eyes and dehydration respectively. The surgical area was cleaned with ethanol and sanitized with an iodine solution (Vetedine). Lidocaine was administered subcutaneously at the incision site. Mice were placed on a stereotaxic apparatus and a craniotomy was performed over the area of interest. Saline solution was regularly applied to the skull.

Injection pipette was lowered to the APT coordinates (Bregma: −2.70 to −2.80mm; Mediolateral: ± 1.10 to 1.15mm; Depth: −2.65 to −2.70mm from the dura) and either, 1μL of AAV8-hSyn-mCherry-Cre (4.9×10^12^ppm) or 1μL of AAV8-hSyn-mCherry (4.6×10^12^ppm) were injected bilaterally with a graduated injection wheel (Narishige, 100μL/rev) at a rate of 0.1 to 0.2μL/min. 5 to 10 min after injection, the pipette was slowly raised. The wound edges were then put back in place and the skin sutured with Vicryl 4-0 thread (Ethicon) or surgical glue (Vetbond 3M). Mice were kept on a heating pad until recovery from anesthesia before being returned to their home cages.

#### Behavioral tests

Two series of behavioral tests were conducted separately in two different laboratories by different experimenters and are presented independently in Figure 4 and supplementary Figure 4. Mechanical and cold sensitivity was tested in preventively injected KI and KO mice, as well as in curatively injected KI and KO SNI mice in Clermont-Ferrand. Mechanical sensitivity in preventively injected KI and KO mice was also assessed in Paris, along with locomotor tests. Both series of experiments yielded similar results, despite notable differences in the paw withdrawal thresholds measured, which can be explained by the differences of experimental conditions.

##### Mechanical sensitivity

Mechanical sensitivity was assessed using Von Frey method. Mice were habituated to the testing environment before baseline testing. On the day of behavior testing, mice were placed in compartments on an elevated grid to allow access to the paw. Different filaments, ranging from 0.02 to 1.4 g, were applied perpendicularly to the plantar surface of the operated paw. 50% paw withdrawal threshold (PWT) was determined using an adaptation of the up and down method (Chaplan et al. 1994).

##### Cold sensitivity

Cold sensitivity was assessed by immersing the operated paw in a 18°C water bath until withdrawal or shaking was observed. In order to avoid stressing mice that are manually tethered in a piece of tissue, they were habituated to the test for 3 days before the baseline test in room temperature water. The first two latencies measured with a difference of less than 2 seconds were averaged to obtain the pain withdrawal latency.

##### Motor behaviors

Motor coordination assessments were performed using the rotarod test. Mice were placed on the rotarod device (Bioseb) during 3 minutes with an accelerating ramp of 4 to 6 rounds per minute. The latency to fall was automatically measured. The test was performed three times for each day of measurement, with two baseline measurements and one measurement at the end of the neuropathic tests. Spontaneous locomotion was also estimated at the end of the neuropathic tests using the circular corridor test. Mice were placed in the cyclotron (IMetronic) in the dark. Four detectors located around the corridor allowed the measurement of the number of quarter turns performed by the animals during 2 hours.

### QUANTIFICATION AND STATISTICAL ANALYSIS

No statistical methods were used to predetermine sample sizes, which are comparable to many studies using similar techniques and animal models.

#### Imaging data analysis

Confocal images acquired for the evaluation of co-expression of Cav3.2-GFP, PV and NeuN were processed using the Fiji/ImageJ software, with the Cell Counter plug-in for manual cell counts. Quantification of the co-labeling of GFP and NeuN were performed on 3 mice (2males, 1 female) by analyzing 11 to 18 images in 5 to 8 brain slices per mouse. Quantification of the co-labeling of GFP and PV were performed on 3 mice (2males, 1 female) by analyzing 10 to 12 images in 5 to 8 brain slices per mouse. Data are expressed as mean ± standard deviation.

#### Electrophysiological data analysis

For in vivo data analysis, potential single-units were first identified using automated clustering software utilizing peak and trough feature sets (KlustaKwik). These clusters were then examined manually for waveform shape (SpikeSort3D, Neuralynx). Upon examination of the interspike intervals, multi-unit clusters were discarded.

Bursts consisted of an initial spike that was followed by one or more spikes at an interval of equal to or less than 5 ms. Since we were interested in potentially T-type channel-mediated bursts, another exclusion criterion was added: bursts had to be preceded by a pause in the spiking activity of at least 50 ms.

Quantification and statistical analysis of in vivo and in vitro data were performed with the Igor Pro v6 and Matlab 2019b softwares, respectively. Between conditions comparison was based on Wilcoxon rank-sum test (Mann-Whitney U test) and Wilcoxon signed rank test for unpaired and paired datasets, respectively. Differences were considered significant if the P-value was lower than 0.05. All data were presented as the means ± standard deviation.

#### Behavioral data analysis

Behavioral data analysis and statistics were performed in R (version 4.1.0), using the RStudio software. Data normality was checked using the Shapiro-Wilks test and no normal distribution was found throughout the datasets. For mechanical and cold sensitivity, overall intra-group comparisons were performed using the Friedman test. Significant differences were identified using a paired Wilcoxon rank-sum test with Bonferroni correction. As no significant difference was found between baseline measurements, each neuropathic measurement was compared to global baseline measurements with the same statistical test. Global inter-group comparisons (KI vs KO) were performed using the Kruskal-Wallis test. Significant differences were identified using the unpaired Wilcoxon rank-sum test. Differences were considered significant if the p-value was lower than 0.05. All values are expressed as mean ± standard error to the mean.

### KEY RESOURCES TABLE

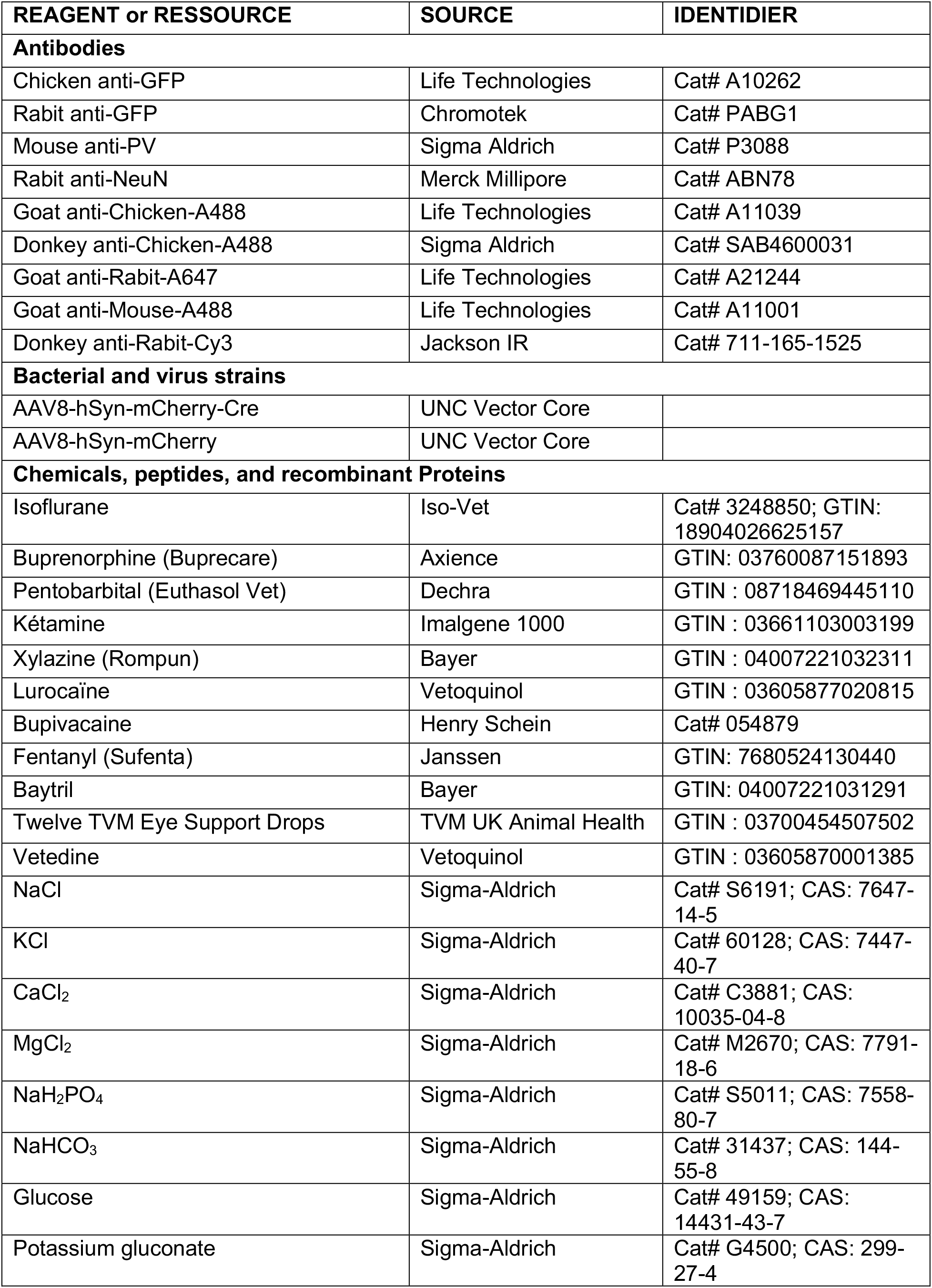
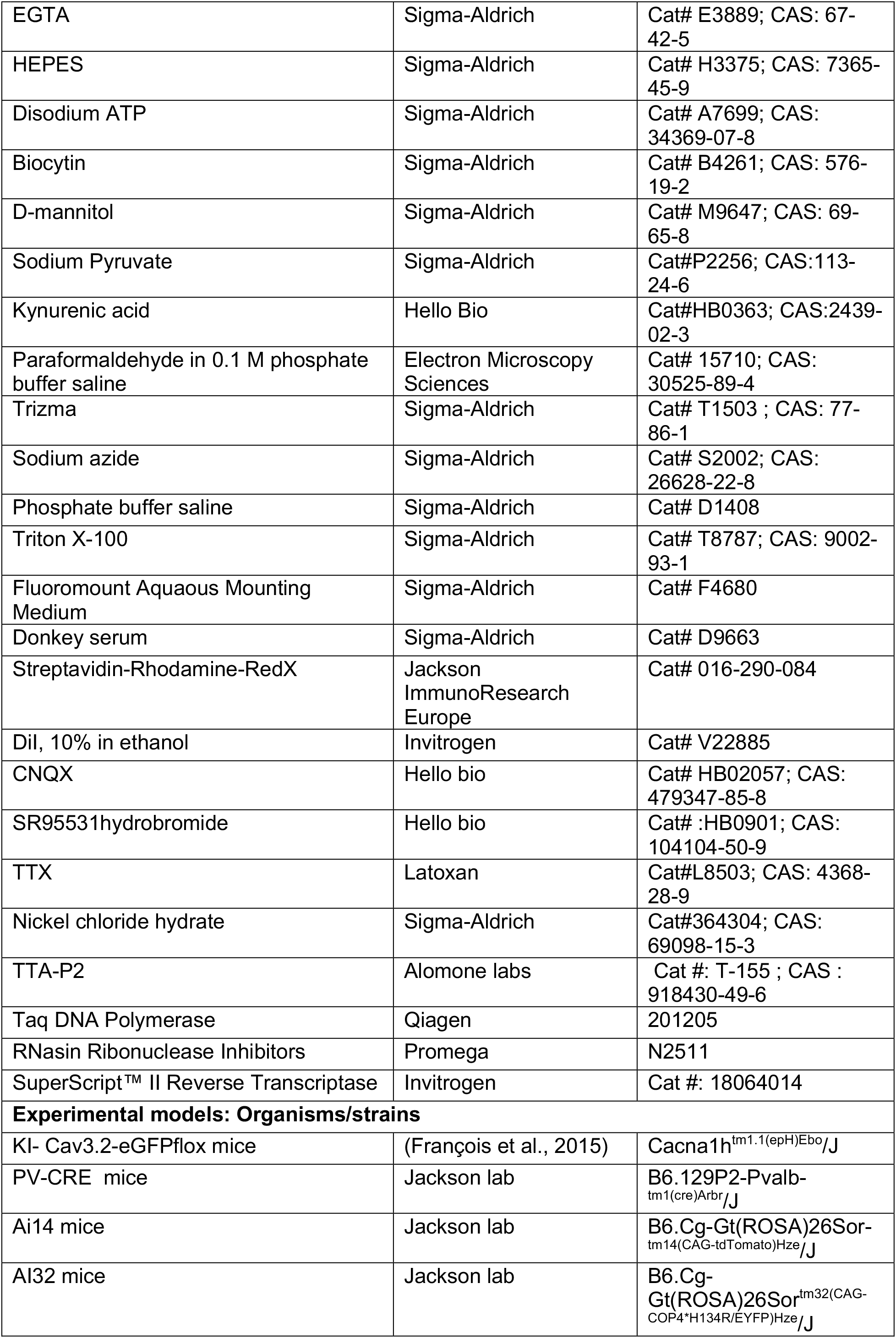
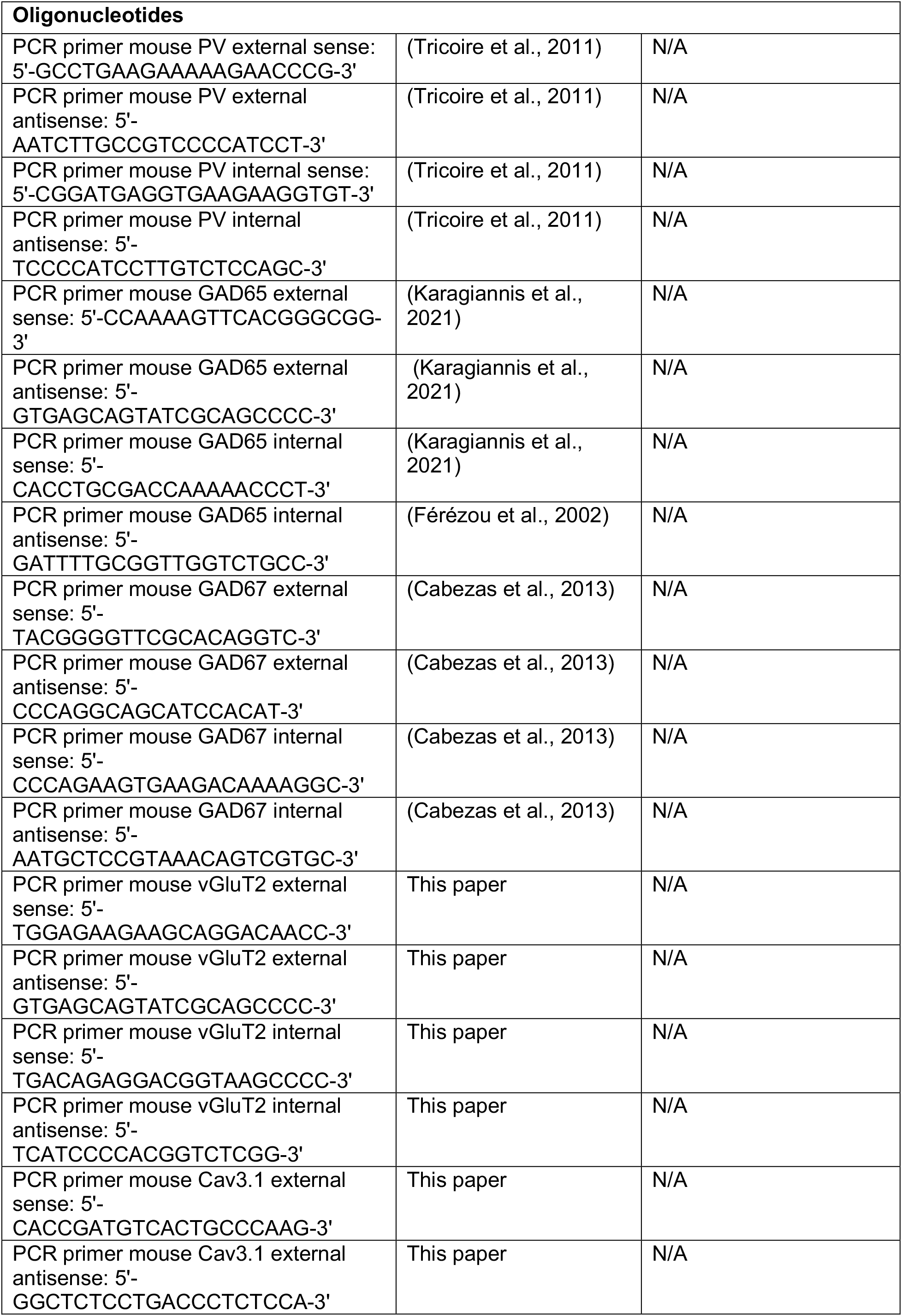
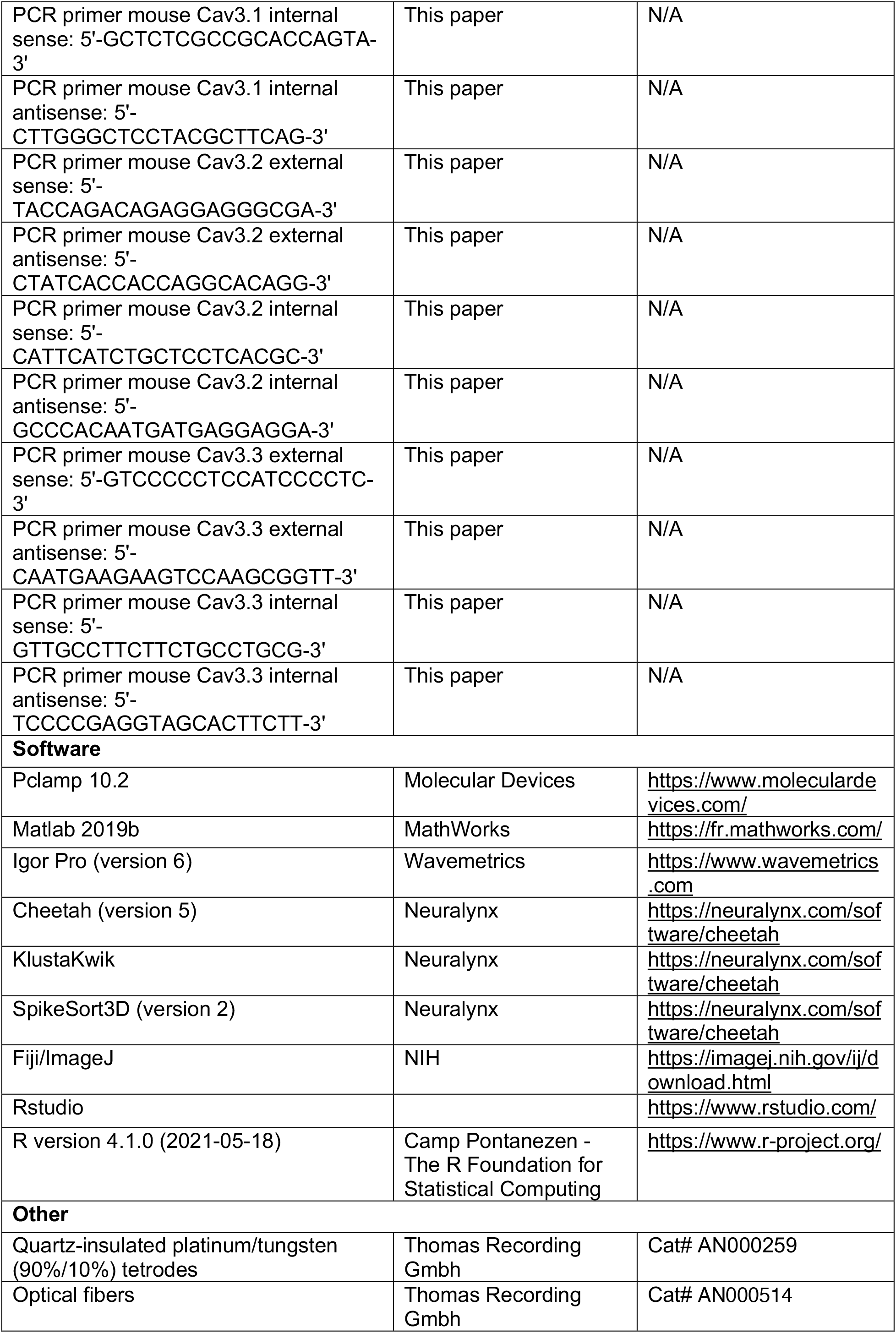
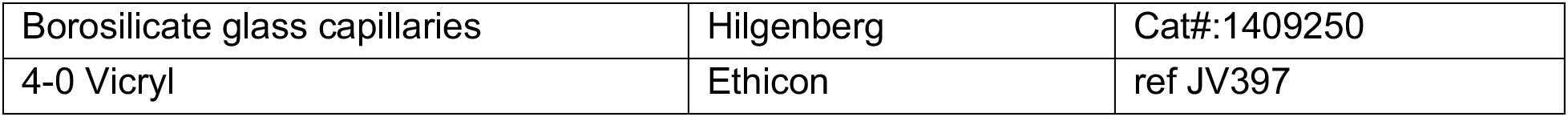

## References

– Bennett, D.L.H., and Woods, C.G. (2014). Painful and painless channelopathies. The Lancet Neurology 13, 587–599.

– Bokor, H., Frère, S.G.A., Eyre, M.D., Slézia, A., Ulbert, I., Lüthi, A., and Acsády, L. (2005). Selective GABAergic Control of Higher-Order Thalamic Relays. Neuron 45, 929–940.

– Bourinet, E., Alloui, A., Monteil, A., Barrère, C., Couette, B., Poirot, O., Pages, A., McRory, J., Snutch, T.P., Eschalier, A., et al. (2005). Silencing of the Cav3.2 T-type calcium channel gene in sensory neurons demonstrates its major role in nociception. EMBO J 24, 315–324.

– Bourinet, E., Francois, A., and Laffray, S. (2016). T-type calcium channels in neuropathic pain. 157, 8.

– Cabezas, C., Irinopoulou, T., Cauli, B., and Poncer, J.C. (2013). Molecular and functional characterization of GAD67-expressing, newborn granule cells in mouse dentate gyrus. Front Neural Circuits 7, 60.

– Candelas, M., Reynders, A., Arango-Lievano, M., Neumayer, C., Fruquière, A., Demes, E., Hamid, J., Lemmers, C., Bernat, C., Monteil, A., et al. (2019). Cav3.2 T-type calcium channels shape electrical firing in mouse Lamina II neurons. Sci Rep 9, 3112.

– Cauli, B., Audinat, E., Lambolez, B., Angulo, M.C., Ropert, N., Tsuzuki, K., Hestrin, S., and Rossier, J. (1997). Molecular and Physiological Diversity of Cortical Nonpyramidal Cells. J. Neurosci. 17, 3894–3906.

– Chaplan, S.R., Bach, F.W., Pogrel, J.W., Chung, J.M., and Yaksh, T.L. (1994). Quantitative assessment of tactile allodynia in the rat paw. Journal of Neuroscience Methods 53, 55–63.

– Chen, W-K., Liu, I.Y., Chang, Y.-T., Chen, Y.-C., Chen, C.-C., Yen, C.-T., Shin, H.-S., and Chen, C.-C. (2010). Cav3.2 T-Type Ca2+ Channel-Dependent Activation of ERK in Paraventricular Thalamus Modulates Acid-Induced Chronic Muscle Pain. J. Neurosci. 30, 10360–10368.

– Decosterd, I., and Woolf, C.J. (2000). Spared nerve injury: an animal model of persistent peripheral neuropathic pain. Pain 87, 149–158.

– Devienne, G., Le Gac, B., Piquet, J., and Cauli, B. (2018). Single Cell Multiplex Reverse Transcription Polymerase Chain Reaction After Patch-clamp. J Vis Exp 57627.

– Dumenieu, M., Senkov, O., Mironov, A., Bourinet, E., Kreutz, M.R., Dityatev, A., Heine, M., Bikbaev, A., and Lopez-Rojas, J. (2018). The Low-Threshold Calcium Channel Cav3.2 Mediates Burst Firing of Mature Dentate Granule Cells. Cereb Cortex 28, 2594–2609.

– Férézou, I., Cauli, B., Hill, E.L., Rossier, J., Hamel, E., and Lambolez, B. (2002). 5-HT3 Receptors Mediate Serotonergic Fast Synaptic Excitation of Neocortical Vasoactive Intestinal Peptide/Cholecystokinin Interneurons. J Neurosci 22, 7389–7397.

– François, A., Schüetter, N., Laffray, S., Sanguesa, J., Pizzoccaro, A., Dubel, S., Mantilleri, A., Nargeot, J., Noël, J., Wood, J.N., et al. (2015). The Low-Threshold Calcium Channel Cav3.2 Determines Low-Threshold Mechanoreceptor Function. Cell Reports 10, 370–382.

– Genaro, K., Fabris, D., and Prado, W.A. (2019). The antinociceptive effect of anterior pretectal nucleus stimulation is mediated by distinct neurotransmitter mechanisms in descending pain pathways. Brain Research Bulletin 146, 164–170.

– Gerke, M.B., Duggan, A.W., Xu, L., and Siddall, P.J. (2003). Thalamic neuronal activity in rats with mechanical allodynia following contusive spinal cord injury. Neuroscience 117, 715–722.

– Giber, K., Slézia, A., Bokor, H., Bodor, Á.L., Ludányi, A., Katona, I., and Acsády, L. (2008). Heterogeneous output pathways link the anterior pretectal nucleus with the zona incerta and the thalamus in rat. J. Comp. Neurol. 506, 122–140.

– Jagodic, M.M., Pathirathna, S., Joksovic, P.M., Lee, W., Nelson, M.T., Naik, A.K., Su, P., Jevtovic-Todorovic, V., and Todorovic, S.M. (2008). Upregulation of the T-Type Calcium Current in Small Rat Sensory Neurons After Chronic Constrictive Injury of the Sciatic Nerve. Journal of Neurophysiology 99, 3151–3156.

– Jeanmonod, D., Magnin, M., and Morel, A. (1996). Low–threshold calcium spike bursts in the human thalamus: Common physiopathology for sensory, motor and limbic positive symptoms. Brain 119, 363–375.

– Jones, A.F., and Sheets, P.L. (2020). Sex-Specific Disruption of Distinct mPFC Inhibitory Neurons in Spared-Nerve Injury Model of Neuropathic Pain. Cell Reports 31, 107729.

– Karagiannis, A., Gallopin, T., Lacroix, A., Plaisier, F., Piquet, J., Geoffroy, H., Hepp, R., Naudé, J., Le Gac, B., Egger, R., et al. (2021). Lactate is an energy substrate for rodent cortical neurons and enhances their firing activity. ELife 10, e71424.

– Kerckhove, N., Mallet, C., François, A., Boudes, M., Chemin, J., Voets, T., Bourinet, E., Alloui, A., and Eschalier, A. (2014). Cav3.2 calcium channels: The key protagonist in the supraspinal effect of paracetamol. PAIN 155, 764–772.

– Kim, D., Song, I., Keum, S., Lee, T., Jeong, M.-J., Kim, S.-S., McEnery, M.W., and Shin, H.-S. (2001). Lack of the Burst Firing of Thalamocortical Relay Neurons and Resistance to Absence Seizures in Mice Lacking α1G T-Type Ca2+ Channels. Neuron 31, 35–45.

– Kim, D., Park, D., Choi, S., Lee, S., Sun, M., Kim, C., and Shin, H.-S. (2003). Thalamic Control of Visceral Nociception Mediated by T-Type Ca2+Channels. Science 302, 117–119.

– Lambert, R.C., Bessaïh, T., Crunelli, V., and Leresche, N. (2014). The many faces of T-type calcium channels. Pflugers Arch. 466, 415–423.

– Lambolez, B., Audinat, E., Bochet, P., Crépel, F., and Rossier, J. (1992). AMPA receptor subunits expressed by single Purkinje cells. Neuron 9, 247–258.

– Lee, J.-H., Gomora, J.C., Cribbs, L.L., and Perez-Reyes, E. (1999). Nickel Block of Three Cloned T-Type Calcium Channels: Low Concentrations Selectively Block α1H. Biophysical Journal 77, 3034–3042.

– Lee, S.E., Lee, J., Latchoumane, C., Lee, B., Oh, S.J., Saud, Z.A., Park, C., Sun, N., Cheong, E., Chen, C.C., et al. (2014). Rebound burst firing in the reticular thalamus is not essential for pharmacological absence seizures in mice. Proc. Natl. Acad. Sci. U.S.A. 111, 11828–11833.

– Lee J-H, Gomota JC, Cribbs LL, and Perez-Reyes E (1999). Nickel Block of Three Cloned T-Type Calcium Channels: Low Concentrations Selectively Block a1H | Elsevier Enhanced Reader.

– Lenz, F.A., Kwan, H.C., Dostrovsky, J.O., and Tasker, R.R. (1989). Characteristics of the bursting pattern of action potentials that occurs in the thalamus of patients with central pain. Brain Research 496, 357–360.

– Lima, S.Q., Hromádka, T., Znamenskiy, P., and Zador, A.M. (2009). PINP: A New Method of Tagging Neuronal Populations for Identification during In Vivo Electrophysiological Recording. PLoS One 4, e6099.

– Llinas, R.R., Ribary, U., Jeanmonod, D., Kronberg, E., and Mitra, P.P. (1999). Thalamocortical dysrhythmia: A neurological and neuropsychiatric syndrome characterized by magnetoencephalography. Proceedings of the National Academy of Sciences 96, 15222–15227.

– Messinger, R.B., Naik, A.K., Jagodic, M.M., Nelson, M.T., Lee, W.Y., Choe, W.J., Orestes, P., Latham, J.R., Todorovic, S.M., and Jevtovic-Todorovic, V. (2009). In vivo silencing of the CaV3.2 T-type calcium channels in sensory neurons alleviates hyperalgesia in rats with streptozocin-induced diabetic neuropathy. Pain 145, 184–195.

– Murray, P.D., Masri, R., and Keller, A. (2010). Abnormal Anterior Pretectal Nucleus Activity Contributes to Central Pain Syndrome. J. Neurophysiol. 103, 3044–3053.

– Pellegrini, C., Lecci, S., Lüthi, A., and Astori, S. (2016). Suppression of Sleep Spindle Rhythmogenesis in Mice with Deletion of CaV3.2 and CaV3.3 T-type Ca(2+) Channels. Sleep 39, 875–885.

– Prado, W.A., and Roberts, M.H.T. (1985). An assessment of the antinociceptive and aversive effects of stimulating identified sites in the rat brain. Brain Research 340, 219–228.

– Rees, H., and Roberts, M.H.T. (1993). The anterior pretectal nucleus: a proposed role in sensory processing. PAIN 53, 121–135.

– Rees, H., G, T.M., and T, R.M.H. (1995). Anterior pretectal nucleus facilitation of superficial dorsal horn neurones and modulation of deafferentation pain in the rat. J. Physiol 489, 159–169.

– Roberts, M.H.T., and Rees, H. (1986). The antinociceptive effects of stimulating the pretectal nucleus of the rat. PAIN 25, 83–93.

– Rossaneis, A.C., and Prado, W.A. (2015). The ventral portion of the anterior pretectal nucleus controls descending mechanisms that initiate neuropathic pain in rats. Clin Exp Pharmacol Physiol 42, 704–710.

– Rossaneis, A.C., Genaro, K., Dias, Q.M., Guethe, L.M., Fais, R.S., Del Bel, E.A., and Prado, W.A. (2014). Descending mechanisms activated by the anterior pretectal nucleus initiate but do not maintain neuropathic pain in rats. EJP 19, 1148–1157.

– Takahashi, T., Aoki, Y., Okubo, K., Maeda, Y., Sekiguchi, F., Mitani, K., Nishikawa, H., and Kawabata, A. (2010). Upregulation of Cav3.2 T-type calcium channels targeted by endogenous hydrogen sulfide contributes to maintenance of neuropathic pain. Pain 150, 183–191.

– Talley, E.M., Cribbs, L.L., Lee, J.H., Daud, A., Perez-Reyes, E., and Bayliss, D.A. (1999). Differential distribution of three members of a gene family encoding low voltage-activated (T-type) calcium channels. J. Neurosci. 19, 1895–1911.

– Terenzi, M.G., Rees, H., Morgan, S.J.S., Foster, G.A., and Roberts, M.H.T. (1991). The antinociception evoked by anterior pretectal nucleus stimulation is partially dependent upon ventrolateral medullary neurones. PAIN 47, 231–239.

– Terenzi, M.G., Rees, H., and Roberts, M.H.T. (1992). The pontine parabrachial region mediates some of the descending inhibitory effects of stimulating the anterior pretectal nucleus. Brain Research 594, 205–214.

– Tricoire, L., Pelkey, K.A., Erkkila, B.E., Jeffries, B.W., Yuan, X., and McBain, C.J. (2011). A Blueprint for the Spatiotemporal Origins of Mouse Hippocampal Interneuron Diversity. J Neurosci 31, 10948–10970.

– Villarreal, C.F., Del Bel, E.A., and Prado, A.W. (2003). Involvement of the anterior pretectal nucleus in the control of persistent pain: a behavioral and c-Fos expression study in the rat. Pain 103, 163–174.

– Villarreal, C.F., Kina, V.A.V., and Prado, W.A. (2004). Participation of brainstem nuclei in the pronociceptive effect of lesion or neural block of the anterior pretectal nucleus in a rat model of incisional pain. Neuropharmacology 47, 117–127.

– Weng, H.-R., Lee, J.I., Lenz, F.A., Schwartz, A., Vierck, C., Rowland, L., and Dougherty, P.M. (2000). Functional plasticity in primate somatosensory thalamus following chronic lesion of the ventral lateral spinal cord. Neuroscience 101, 393–401.

